# Inferring cell-type-specific causal gene regulatory networks during human neurogenesis

**DOI:** 10.1101/2022.04.25.488920

**Authors:** Nil Aygün, Dan Liang, Wesley L. Crouse, Gregory R. Keele, Michael I. Love, Jason L. Stein

**Author notes:** These authors jointly supervised the work (Michael I. Love, Jason L. Stein).

## Abstract

**Background:** Genetic variation influences both chromatin accessibility, assessed in chromatin accessibility quantitative trait loci (caQTL) studies, and gene expression, assessed in expression QTL (eQTL) studies. Genetic variants can impact either nearby genes (local eQTLs) or distal genes (trans eQTLs). Colocalization between caQTL and eQTL, or local- and distant-eQTLs suggests that they share causal variants. However, pairwise colocalization between these molecular QTLs does not guarantee a causal relationship. Mediation analysis can be applied to assess the evidence supporting causality versus independence between molecular QTLs. Given that the function of QTLs can be cell-type-specific, we performed mediation analyses to find epigenetic and distal regulatory causal pathways for genes within two major cell types of the developing human cortex, progenitors and neurons.

**Results:** We found that expression of 168 and 38 genes were mediated by chromatin accessibility in progenitors and neurons, respectively. We also found that the expression of 781 and 200 downstream genes were mediated by upstream genes in progenitors and neurons. Moreover, we discovered that a genetic locus associated with inter-individual differences in brain structure showed evidence for mediation of *SLC26A7* through chromatin accessibility, identifying molecular mechanisms of a common variant association to a brain trait.

**Conclusions:** In this study, we identified cell-type-specific causal gene regulatory networks whereby the impacts of variants on gene expression were mediated by chromatin accessibility or distal gene expression. Identification of these causal paths will enable identifying and prioritizing actionable regulatory targets perturbing these key processes during neurodevelopment.

## Background

Genome-wide association studies (GWAS) have identified many common genetic variants associated with risk for neuropsychiatric disorders [1–3] and inter-individual differences in other brain relevant traits, like cortical structure [4–6]. GWAS studies alone do not yield molecular, cellular, and systems level causal pathways by which discovered genetic variation influences a trait. Given the enrichment of brain trait associated variants within non-coding regulatory elements [7, 8], quantitative trait loci (QTL) analyses for gene regulatory phenotypes, including chromatin accessibility and gene expression, have been widely applied for the functional interpretation of GWAS (Fig. 1a) [9–12]. Importantly, recent QTL studies have demonstrated that these non-coding brain trait-associated variants exert their functional effects in a developmental and cell-type-specific manner [13–16].

**Figure 1.**
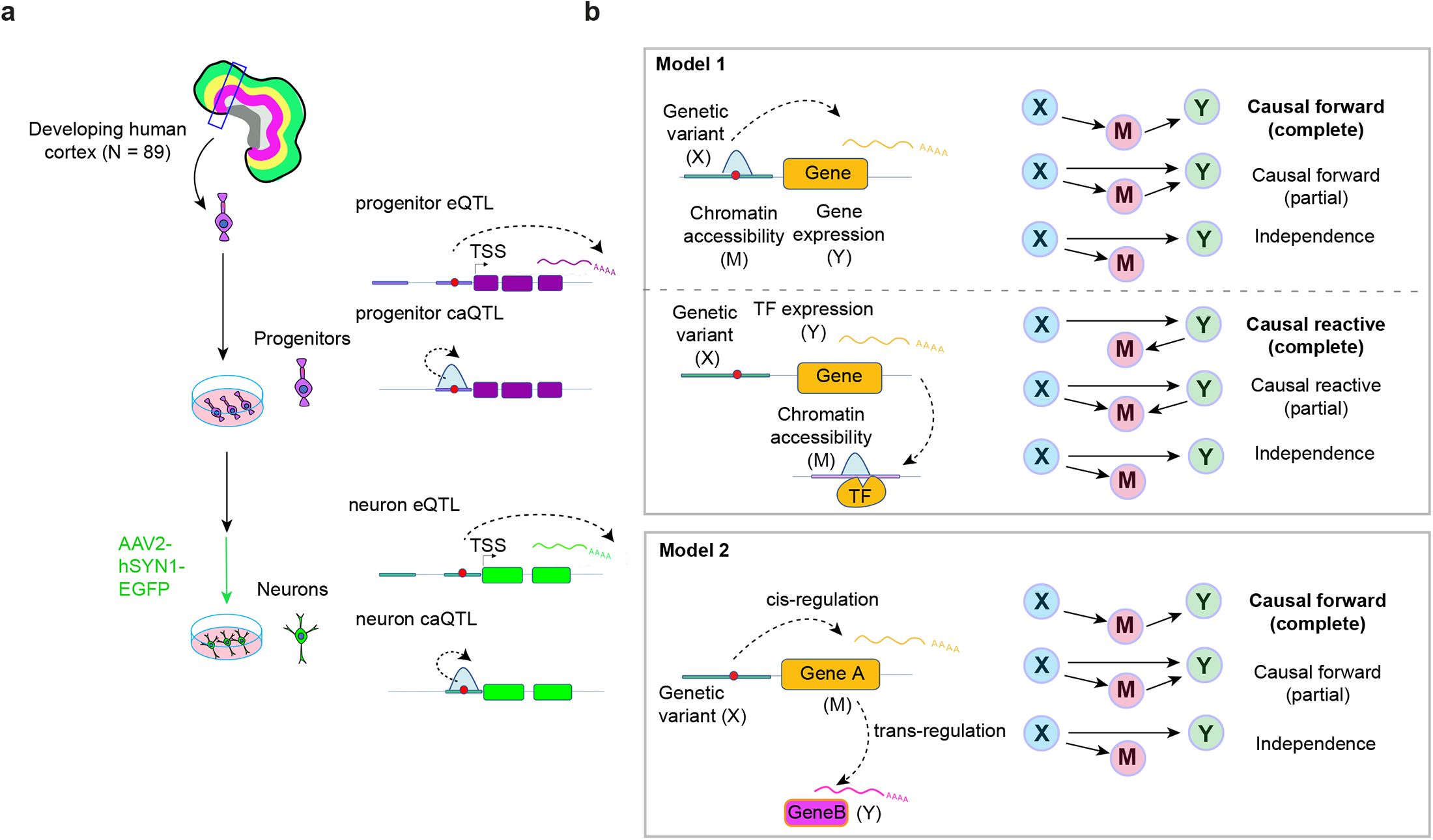
Study design. **a)** Cell-type-specific *in vitro* experimental system including progenitors and their differentiated neuronal progeny. **b)** Models were evaluated in the study.

To find the causal gene regulatory networks by which non-coding genetic variation influences a complex brain or behavioral trait, first a colocalization between a molecular QTL, performed in a relevant cell-type and developmental time period, and the trait GWAS is conducted to identify shared causal variants [11, 17]. Similarly, colocalization between multiple molecular QTLs can be used to determine if there are shared genetic influences on both traits, for example chromatin accessibility and gene expression [18, 19]. However, colocalization does not guarantee a causal relationship. If a genetic variant, represented by X, is significantly associated with both a candidate mediator (M) such as chromatin accessibility, and an outcome (Y) such as gene expression measured from the same individuals; X (or an LD proxy of X) can directly regulate Y independent from M, termed independence (X -> M; X -> Y); the effect of X on Y can be mediated partially or completely through M, termed a forward model (X -> M -> Y); or the effect of X on M can be partially or completely mediated by Y, termed a reactive model (X -> Y -> M) [20–22] (Fig. 1b). The causal ordering of events in each model is not dependent on the directionality of the effect of X on M or Y, which is fixed from X to M or Y. A causal pathway of a brain trait associated variant can be used to prioritize actionable trait relevant therapeutic targets that interrupt the key pathological processes [23–25].

Causal pathways for non-coding trait-associated genetic variation have been experimentally demonstrated for a small subset of GWAS traits. An example of the causal forward model (X -> M - > Y) is a common genetic variant (X) located within a regulatory element (M), such as a gene promoter or enhancer, that disrupts the binding motif of a transcription factor (TF) leading to differential chromatin accessibility observed via chromatin accessibility QTL (caQTL) analysis, which leads to differential expression (Y) observed via cis-expression QTL (cis-eQTL) analysis [7,26–29]. Though less canonical, causal reactive models (X -> Y -> M) have also been experimentally verified, whereby changes in the expression of a gene encoding a TF may cause changes in the chromatin accessibility in the genomic regions harboring a TF binding motif [30, 31]. Similar gene regulatory mechanisms can lead to trans-eQTLs, whereby a genetic variant (X) cis-regulates a transcription factor or components of a signaling cascade (M) and leads to altered expression of a distal gene (Y). Alternatively, in the independence model, the expression of the gene is independent of chromatin accessibility, which may occur due to false positive colocalizations or non-canonical regulatory mechanisms [30,32–36].

There are two common ways to test causal models using genetic association data. Mendelian randomization (MR) analyses infer causal relationships by defining “instruments”, variants that influence M and satisfy certain modeling assumptions, and determining if these instruments may affect Y through M by examining their paired effect sizes and standard errors [37–39]. MR analyses have the advantages of using summary statistics rather than raw data, and additionally two phenotypes M and Y are not required to be measured in the same individuals. However, MR approaches require allelic heterogeneity, which is not compatible with caQTLs and is only detectable with large sample sizes in eQTLs for many genes [12,13,40]. Alternatively, mediation analysis can be performed to distinguish between pleiotropic, forward, and causal reactive models when multiple phenotypes are measured across the same donors and there is access to the raw data. Evidence for the forward model is established in a classical mediation approach when there is no uncontrolled confounding and (1) M and Y are conditionally dependent given X; and (2) X and Y are conditionally independent given M [41, 42]. Mediation approaches have been applied to individual-level multi-modal QTL data to infer causal relationships between different molecular intermediates including mediation of gene expression via DNA methylation, histone acetylation [43], chromatin accessibility [44, 45] and trans-regulation by distal genes [46–50]. However, these previous studies were subject to difficulties with the specification of multiple null hypotheses, correction for multiple testing, and quantifying evidence for complete versus partial mediation. A recent Bayesian model selection framework can overcome these challenges by weighing the evidence of each potential relationship across X-M-Y triplets, summarized as a posterior probability [22].

Current studies have investigated genetically mediated causal gene regulatory networks in the human brain using multi-modal QTLs, but they were limited to studying data derived from bulk adult brain tissue and were not able to resolve cell-type and developmentally specific mechanisms [43,51– 53]. In this study, we applied a Bayesian causal inference method, *bmediatR*, to examine the mediation of genetic effects (1) on gene expression through nearby chromatin accessibility, (2) on distal chromatin accessibility through the expression of TFs and (3) on downstream gene expression through trans-regulation by other genes using our previously generated ca/eQTL datasets derived from human cortical progenitors and their differentiated neuronal progeny (Fig. 1a) [13, 14]. We identified causal paths for gene expression mediated by chromatin accessibility for 168 and 38 genes in progenitors and neurons. Also, we found that the expression of 781 and 200 downstream genes was mediated by upstream genes proximal to cis-regulatory SNPs in progenitors and neurons, respectively. Furthermore, we proposed causal mechanisms affecting brain structure and neuropsychiatric disorders through multiple levels of biology by defining a causal forward regulatory mechanism leading to changes in expression of the *SLC26A7* gene via chromatin accessibility at a locus co-localized with middle temporal gyrus area GWAS [4], and for a gene associated with educational attainment through transcriptome-wide association study (TWAS), *TUBG2* [6], via trans-regulation by *SRP72* gene in progenitors.

## Results

### Cell-type-specific mediation of expression via chromatin accessibility (forward model)

We assessed causal mediation using previously generated cell-type specific ATAC-sequencing [13] and RNA-sequencing [14] data that was subset to a dataset where both data modalities were acquired in the same donors for each cell type (N_donor_ = 75 in progenitors and N_donor_ = 57 in neurons). To identify causal models explaining X-M-Y triplets, we identified variants impacting both chromatin accessibility and gene expression (Fig. 1b, Model 1). We first subset the ca/eQTLs to biologically relevant X-M-Y triplets to test the forward model, as this is the most commonly assumed model to explain genetic effects on both chromatin accessibility and gene expression [7]. The causal forward model was only tested when the ca/eQTL variant (FDR < 5%) was within the chromatin accessible region, because such a variant in a gene regulatory region is likely to disrupt TF binding and then affect gene expression (Fig. 1b, Model 1 upper diagram). We found that in progenitors and neurons, 681 and 204 variants within 289 and 64 chromatin accessible regions +/-1Mb from the transcription start site (TSS) of 332 and 83 genes were significantly associated with both chromatin accessibility and gene expression (Fig. S1a).

In concordance with the previous observation of the sharing of directionality of genetic effect between caQTLs and eQTLs [13, 45], we detected that 85% and 80% of X-M-Y triplets showed allelic effects in the same direction on both chromatin and gene expression progenitor and neurons, respectively (Fig. S1b). To evaluate causal forward relationships of genetic mediation of gene expression through chromatin accessibility, we applied a Bayesian mediation approach *bmediatR* [22] where X is a single variant within the chromatin accessibility peak, M is chromatin accessibility, and Y is the gene expression.

We detected 168 and 38 genes associated with 364 and 87 variants in progenitors and neurons supported by the causal forward model (posterior probability > 0.50; Fig. 2a, Table S1). As an example where the Bayesian approach supported the causal forward model, an eQTL-caQTL colocalization in progenitors showed that variation of *CHL1* gene expression was found to be mediated through a chromatin accessibility peak (chr3:74,521-75,730) (Fig. 2b, posterior probability causal forward complete/causal forward partial: 0.40/0.59). The same progenitor eSNP and caSNP (rs9867864) within a chromatin accessibility peak located 121 kb upstream of the gene TSS was found to influence *CHL1* expression through altering chromatin accessibility, and showed allele-specific chromatin accessibility (ASCA) (FDR for ASCA = 9.5 × 10^−22^). The SNP did not survive our threshold for testing causal models in neurons, because rs9867864 was significantly associated with chromatin accessibility but not with gene expression, indicating that there were cell-type-specific impacts of this variant on gene expression (ca/eQTL nominal p-values in neurons = 1×10^−9^/0.019, Fig. S2a). We also performed a mediation scan analysis substituting different chromatin accessible regions (M) within +/-1MB of the TSS of the gene to test for specificity of the mediation effect (posterior probabilities for substituted regions M are shown in Fig. 2b). This analysis showed that the chromatin accessibility peak containing the variant had the highest posterior probability for forward mediation of the genetic effect on *CHL1* expression (Fig. 2b). Though 94% of peaks tested did not show evidence for mediating the effect, we detected two more chromatin accessibility peaks also surviving the threshold we used to determine causal forward models, although our caQTL analysis was not sensitive enough to detect the significant impact of rs9867864 on these chromatin accessible regions (causal forward complete/causal forward partial for peak1 = 4.8 × 10^−7^/0.69; for peak2 = 1.1 × 10^−6^/0.54, nominal caQTL p-values for peak1 = 0.042, peak2 = 0.085, Fig. 2b). Both peak1 and peak2 were significantly associated with *CHL1* expression after controlling for the center peak and technical covariates (Fig. S2b). This observation suggests that *CHL1* expression may be mediated by the center peak along with peak1 and peak2. There have been case studies, though no definitive associations, suggesting that heterozygous deletion of *CHL1* showed language and cognitive developmental delay [54, 55]. Here, we propose a progenitor-specific causal regulatory mechanism for differences in *CHL1* expression. If the association with CHL1 and cognitive delay is confirmed, this finding may have therapeutic potential in that *CHL1* expression levels could be manipulated through targeting epigenetic engineering tools in progenitors to this enhancer region.

**Figure 2.**
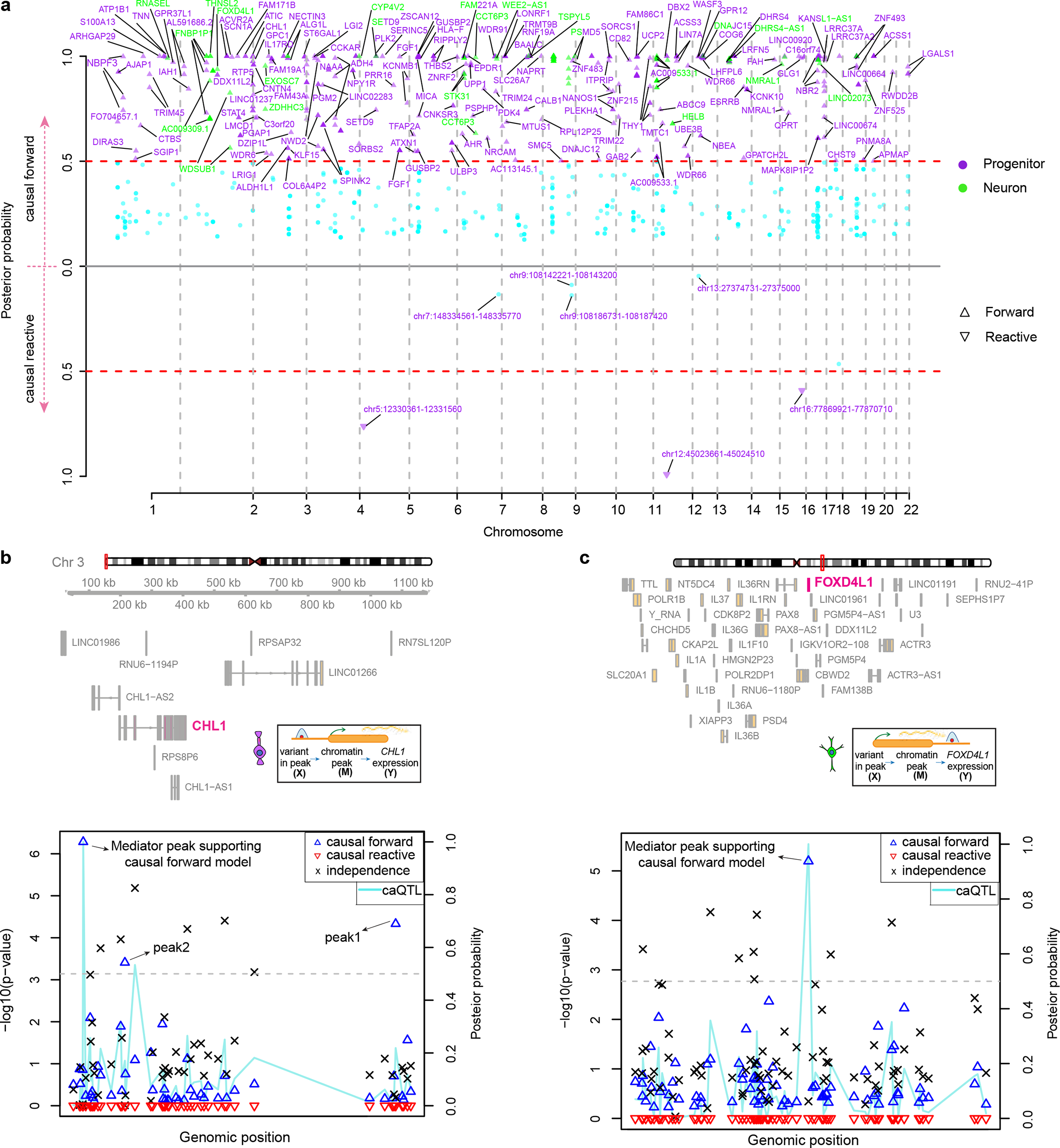
Cell-type-specific gene regulation mediated by chromatin accessibility. **a)** Manhattan plot showing the *bmediatR* results for causal forward model (upper), and for causal reactive model (lower) for each cell-type (purple for progenitor, green for neuron, purple + green for genes detected in both cell-types). Gene symbols and chromatin accessible regions were shown. **b)** Mediation scan plot overlaid with caQTL data for the causal forward model of epigenetic regulation of *CHL1* gene expression in progenitors indicated by lines in different colors with corresponding p-value on the left y-axis, and the posterior probability of each possible model showed with different shapes on the right y-axis. The locations of center peak, peak1 and peak2 supporting causal forward model were highlighted. **c)** Mediation scan plot overlaid with caQTL data for the causal forward model of epigenetic regulation of *FOXD4L1* gene expression in neurons. Line and shapes were assigned as part b.

Another cell-type-specific mediation supporting the causal forward model was observed in neurons at the *FOXD4L1* gene locus. The variant rs141063413, within a chromatin accessible region (chr2:113503031-113503850) located 2kb downstream of the gene was significantly associated with *FOXD4L1* expression and chromatin accessibility in neurons (Fig. 2c). In progenitors, the same variant was associated with chromatin accessibility but not gene expression (ca/eQTL nominal p-values in progenitors = 6.9×10^−10^/0.12, Fig. S2c). The chromatin accessible region including the variant rs141063413 mediated *FOXD4L1* expression in neurons (causal forward complete/causal forward partial: 0.024/0.914, Fig. 2c). FOXD4L1 was shown to be required for embryonic neurogenesis in xenopus [56]. These results again suggest a cell type and a regulatory element that may be useful in modulating the expression of a given gene.

Next, we assessed the features that can be predictive of causal versus independent relationships between chromatin accessibility and gene expression regulated by the same locus. We found that as the relationship between chromatin accessibility and gene expression (percent variance in gene expression explained by chromatin accessibility termed r^2^(Y,M)) is stronger, causal forward models were more strongly supported in both cell types (Fig. 3a). We also observed that e/caQTLs supporting causal forward models were significantly closer to the gene TSS in both cell-types, and enriched more within promoters in progenitors (Fig. 3b-c). However, we did not detect enrichment of testable variants disrupting TF binding motifs [57] nor variants supported by ASCA within the causal forward model compared to the independence model (Fig. 3d-e). This suggests that the amount of variance in gene expression explained by chromatin accessibility and genomic location relative to gene TSS can be predictive features for causality. However, TF motif disruption and ASCA are not reliable features for the assumption of causality. It is important to note that low power to detect ASCA based on allele frequency, incomplete annotation of motifs, and incomplete knowledge of how genetic variation disrupts TF binding, all prevent a comprehensive evaluation of these features.

**Figure 3.**
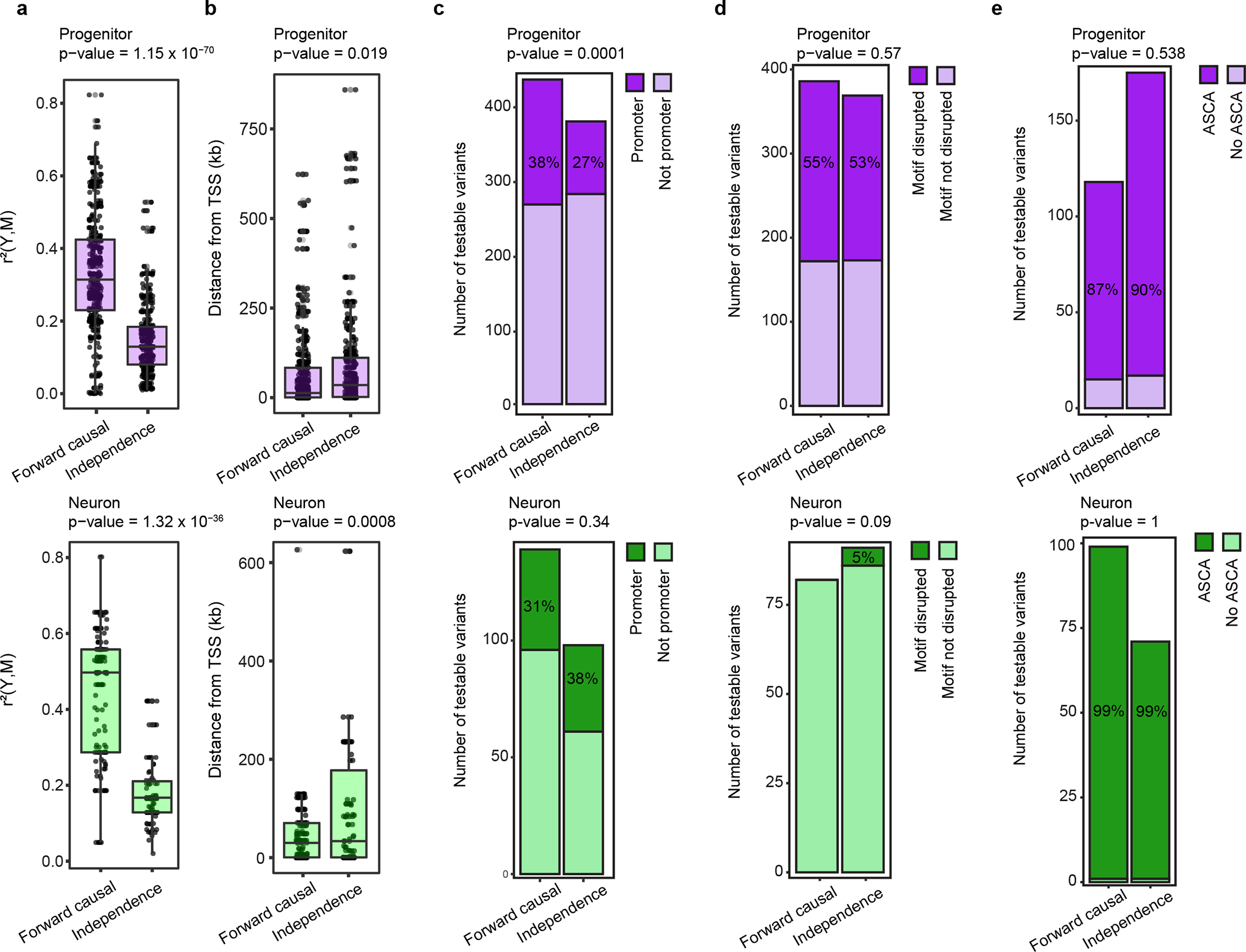
Biological and technical features to predict causality. **a)** Percent variance in gene expression explained by chromatin accessibility (*r^2(^Y,M)*) in X-M-Y triplets supporting causal forward model versus independence in progenitors (upper, purple) and neurons (lower, green). Unpaired t-test p-value was shown. **b)** Absolute distance of variants relative to TSS of the genes for X-M-Y triplets supporting causal forward model versus independence in progenitors (upper, purple) and neurons (lower, green). Unpaired t-test p-value was shown. **c)** Distribution of variants located within gene promoter (within +/-2kb window from gene TSS) within X-M-Y triplets supporting causal forward model versus independence in progenitors (upper, purple) and neurons (lower, green). Chi-square test p-value was shown. **d)** Number of testable variants detected to disrupt TF binding motifs via *motifbreakR* in X-M-Y triplets supporting causal forward model versus independence in progenitors (upper, purple) and neurons (lower, green). Chi-square test p-value was shown. **e)** Number of testable variants (either variant itself or LD proxy) showed allele-specific chromatin accessibility (ASCA) in X-M-Y triplets supporting causal forward model versus independence in progenitors (upper, purple) and neurons (lower, green). Chi-square test p-value was shown.

A comparison of triplets supporting the causal forward model using the Bayesian approach (bmediatR) with regression-based mediation analysis [44] showed that 90.7% and 23.7% of the triplets in progenitors and neurons overlapped with the triplets supporting causal forward models determined by a complementary regression-based approach (Fig. S3a). We hypothesized that lower statistical power may at least partially explain the lower agreement between the two approaches in neurons. To this end, we down-sampled progenitors to be equal in sample size with neurons, and as expected we found a decreased overlap between Bayesian versus regression-based mediation approaches (Fig. S3b, 47.5% overlap). This suggests that there may be general agreement between the two approaches for studies with a sufficient sample size.

### Accuracy of Bayesian mediation approach in classifying forward versus reactive models given differences in measurement error between phenotypes

Given that differences in measurement error between candidate mediators and outcomes may give rise to false positive causal relationships [20, 23], we quantified measurement error using the intraclass correlation coefficient (ICC) [58] across technical replicates from the same donor line thawed multiple times. We observed that ICC for ATAC-seq measured peaks (Ndonors_with_replicates = 11 in progenitors, Ndonors_with_replicates = 5 in neurons) were on average lower than RNA-seq measured genes (Ndonors_with_replicates = 13 in progenitors, Ndonors_with_replicates = 9 in neurons) (Fig. S4a), indicating higher measurement error in ATAC-seq data. We simulated the impact of measurement error on causal models to find the ICC at which true causal forward or reactive models flip to being falsely called reactive or forward models. Since we observed that the ICC flipping threshold was also dependent on the magnitude of the effects of X on M and Y on M, we calculated an ICC threshold by varying the magnitude of effects for X-M-Y triplets. For example, in a simulated causal forward model where a variant X explains 30% of the variation in chromatin accessibility (M) and the variant X also explains 10% of the variation in gene expression (Y), we find that ICC values of chromatin accessibility below 0.3 lead to incorrect model flipping (Fig. S4b-c; see Methods). We filtered out the triplets supporting causality with ICC values lower than per triple threshold of ICC at which model flipping occurred for all future analyses (Fig. S4b, see Methods).

We then subset the ca/eQTLs to biologically relevant X-M-Y triplets to test the reactive models. The causal reactive model was only tested when the ca/eQTL variant altered the expression of a TF and the chromatin accessibility of a region harboring a motif for that TF, as this is a likely mechanism for a causal reactive model (Fig. 1b, Model 1 lower diagram). We detected 3 variants cis-regulating 3 genes encoding TFs and also associated with chromatin accessibility within 8 regions that contained the binding motif of the TF in progenitors (Fig. 2a). We did not detect any significant ca/eQTLs matching our causal reactive model criteria in neurons (at 5% FDR for trans-caQTLs).

We would expect that in true reactive models, where a variant’s effect on chromatin accessibility is fully mediated through gene expression, no allelic imbalance in chromatin accessibility would be observed by X or an LD proxy of X, because the effect is not *cis* with respect to the chromatin peak, but *trans*. Hence, we excluded variants within chromatin accessible regions to evaluate causal reactive models. We investigated a few cases where the reactive model had a higher posterior probability than the forward model. As an example, the variant rs2731040 located 1 kb upstream, not within, a chromatin accessible region (chr12:45023661-45024510) harboring several DBX2 TF binding motif sites was significantly associated with both *DBX2* expression and chromatin accessibility (Fig. S5a-b). This example satisfied our previously defined biologically motivated criteria to test whether the data support a causal reactive model. The same chromatin accessible region additionally harbored binding motifs of 15 different TFs, the expression of each of which was cis-regulated. When we tested if the variants cis-regulating expression of these TFs were also significantly associated with chromatin accessibility at this peak, we did not detect any significant trans-caQTL loci other than rs2731040, the variant that was also associated with *DBX2* TF expression (Fig. S5c). Data from this X-M-Y triplet best fit the causal reactive model (posterior probability for causal forward/causal reactive: 0.0095/0.99). This observation initially suggested a potential autoregulatory mechanism for a reactive model whereby expression of *DBX2* regulated by rs2731040 variant subsequently led to a change in chromatin accessibility. However, we also found another variant, rs2731038, within the chromatin accessible region that was in LD with rs2731040 (r^2^ = 0.7), a variant that showed ASCA in our previous study (Fig. S5d, the adjusted p-value for ASCA = 0.02) [13]. Given that ASCA indicates a forward regulatory mechanism for chromatin accessibility by the variant, we interpret the high posterior probability supporting the reactive model as a false positive result. This observation shows that causal reactive models must be carefully evaluated to ensure that they are not tagging allele-specific effects, which are indicative of causal forward models. Through careful follow-up of X-M-Y triplets, we demonstrate that false positive reactive models may persist despite stringent criteria and posterior probability thresholding.

### Cell-type-specific mediation of eQTLs via trans-regulation

Next, we investigated genes that were mediated via trans-regulation for each cell-type using the same Bayesian mediation approach. Because this analysis did not require both ca/eQTL data derived from the same donor, sample sizes were slightly increased as compared to the previous analyses (N_donor_ = 85 in progenitors and N_donor_ = 74 in neurons) (Fig. 1b, Model 2). Within each cell-type, SNPs with significant association to a proximal gene (cis-eSNPs) were also tested for association with every other gene (trans-eQTL, significance threshold p-value < 1×10^−6^ was applied, consistent with previous trans-eQTL studies [46, 47]). Among those trans-eQTLs surviving the significance threshold, we discovered 781 and 200 downstream genes that were mediated via trans-regulation by 225 and 107 cis-regulated upstream genes in progenitors/neurons, respectively (Fig. 4a; posterior probability for causal forward > 0.5, Table S2). Sample size differences between the cell-types were likely a large contributor to the differences in the number of mediated trans-QTLs detected. We additionally used an alternative regression-based mediation approach called MOSTWAS [47] and found that 22.3% and 10% of trans-regulated (downstream) genes detected via the Bayesian mediation approach in progenitor and neurons, respectively, were also supported by this alternative approach (Fig. S6).

**Figure 4.**
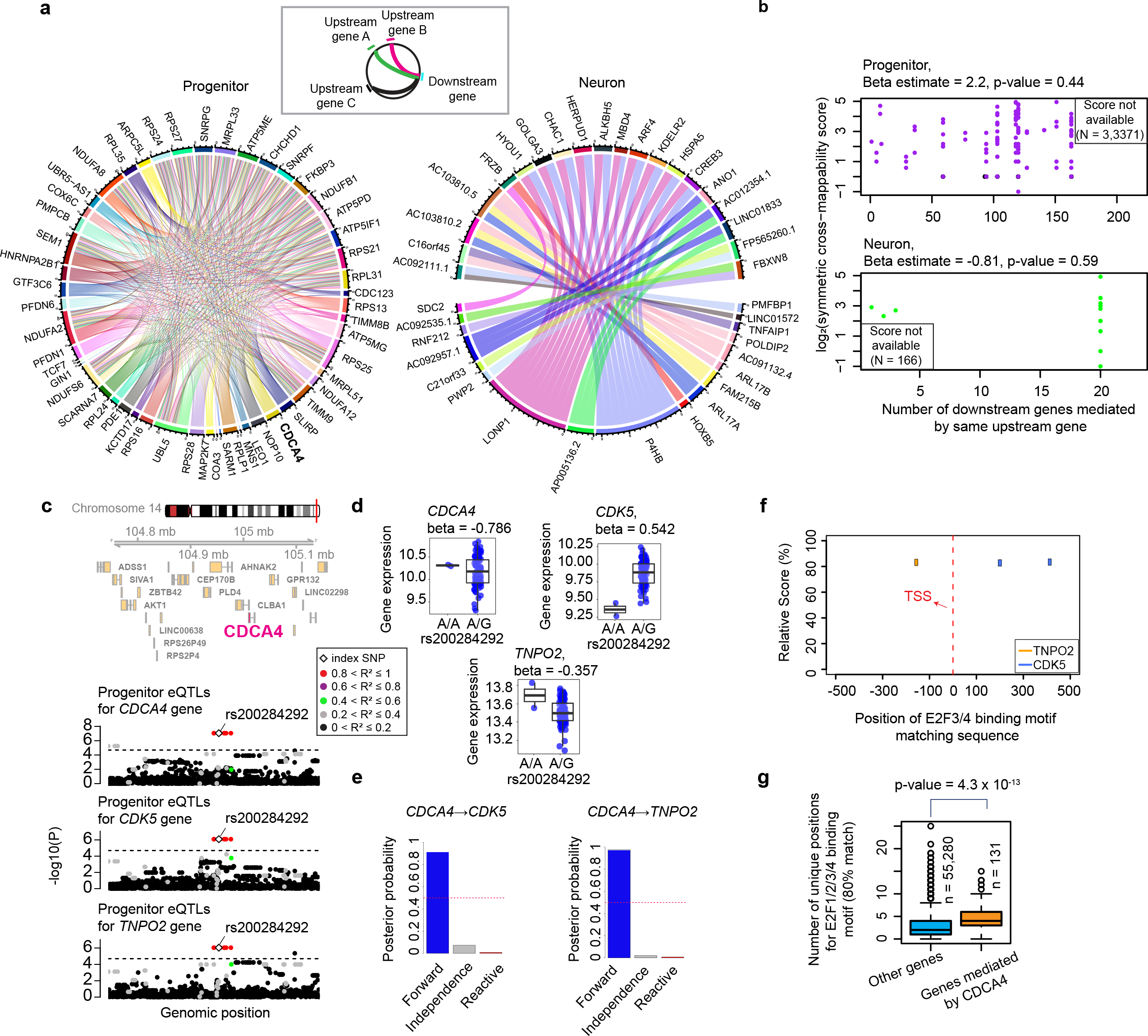
Cell-type-specific gene expression mediated by trans-regulation. **a)** Circle plots illustrating a subset of genes (top 20 downstream genes ranked by the number of upstream genes regulating them) mediated by trans-regulation in progenitors at left and in neurons at the right panel. Each upstream gene was shown as different colored lines interacting with different downstream genes. **b)** The relationship with symmetric cross mappability scores on the y-axis with the number of downstream genes mediated by the same upstream gene in progenitors (upper panel) and neurons (lower panel). Number of upstream-downstream gene pairs for which cross-mappability scores were not available were shown. **c)** Genomics tracks illustrating association of variants cis-regulating *CDCA4* upstream gene with the expression *CDCA4*, and downstream genes *CDK5* and *TNPO2*. Data points were colored based on the pairwise LD r^2^ with the rs200284292. **d)** Boxplots showing the relationship between expression of genes and rs200284292 variant. **e)** bmediatR posterior probability for causal forward, independence versus causal reactive models for the regulation of C*DK5* and *TNPO2* genes by *CDCA4*. **f)** C*DK5* and *TNPO2* gene loci within +/-500 bp of gene TSS showing the genomic regions matching the binding motif of E2F3 and E2F4 transcription factors at the 80% of matching rate. **g)** Enrichment of E2F1, E2F2, E2F3 and E2F4 TF matching motifs within the promoters of downstream genes of *CDCA4* versus the rest of genes in the genome.

We found that cell-type-specific cis-regulated genes (upstream genes) mediated the expression of multiple trans-genes (downstream genes). For example, upstream genes *CDCA4* and *LONP1* mediated the expression of 123 and 20 downstream genes in progenitors and neurons, respectively (Fig. 4b). High cross-mappability score, a statistic quantifying sequence similarity between two genes, has been shown to often result in false positive trans-eQTL discovery [59]; therefore, we obtained the cross-mappability score of the upstream genes that mediate multiple downstream genes. We did not observe cross-mappability as a major driving factor for the number of cis-genes involved in trans-regulation (Fig. 4b).

As an example, *CDCA4* on chromosome 14, was cis-regulated by a progenitor specific eSNP (rs200284292) and mediated the expression of genes *CDK5* on chromosome 7 and *TNPO2* on chromosome 19 (Fig. 4c and 4d, posterior probabilities supporting causal forward = 0.874 and 0.958 for *CDK5* and *TNPO2,* Fig. 4e). CDCA4 has been found to repress E2F transcription factor family-dependent transcriptional activation [60], and ChIP-seq data in another study showed binding sites of E2F4 and E2F3 within *CDK5* and *TNPO2* gene promoters, respectively, in the mouse neural precursor cells [61]. Consistent with CDCA4 influencing expression of downstream genes via E2F family TFs, we detected multiple matching sequences for binding motifs of E2F4 and E2F3 TFs within promoters of both *CDK5* and *TNPO2* genes (Fig. 4f). We also found an increase in E2F1-4 TF motifs within the promoter of 131 genes mediated by *CDCA4* gene as compared to all other protein-coding genes which did not show evidence of mediation (Fig. 4g). Loss-of-function mutations in CDK5 protein were previously associated with lissencephaly [62], and *de novo* mutations in TNPO2 protein were found within individuals with neurodevelopmental abnormalities [63]. *CDCA4* may therefore represent an important master regulator of disease associated genes in progenitors during neural development.

We also found support for one trans-gene being causally mediated by multiple cis-genes. This occurred in ribosomal genes including *RPS27* (Nprogenitor-upstream-gene = 31) and *RPL11* (Nprogenitor-upstream-gene = 26) (Fig. 4a). This finding supports the previous observation that trans-regulation highly influences the expression of ribosomal genes via potential miRNA and lncRNA actions [64].

In this analysis, we tested all cis-eQTL variants regardless of LD for trans-eQTL causal analyses. We observed that only 1.2% and 0.24% of variants in progenitor and neurons which involved in trans-regulation were conditionally-independent index variants discovered in cis-eQTL analysis for the upstream genes. This may suggest that performing mediation analysis for trans-eQTL discovery enables the detection of additional genomic loci that were disregarded since they were conditionally dependent on index variants; however, they could play an instrumental role in regulating gene expression.

### Proposing regulatory mechanisms of brain-related GWAS loci via genetically mediated gene expression

We further leveraged cell-type-specific causal pathways to interpret the function of GWAS loci associated with brain-relevant traits (Table S3). As a specific example, we observed that an indel variant (rs10717382) significantly associated with chromatin accessibility (chr8:91179881-91181040) at the promoter of the *SLC26A7* gene as well as its expression in progenitors was also co-localized with an index SNP (rs57117164) associated with inter-individual differences in the surface area of a specific cortical region, the Middle Temporal Gyrus [4] (Fig. 5a). Importantly, we detected that deletion of the T allele decreased the binding affinity of the NKX2-2 transcription factor based on *in silico* analysis (Fig. 5b) [57]. NKX2-2 was previously found to be functioning as a transcriptional repressor [65–67], and consistent with this, we observed that the deletion of the T allele was also associated with increased chromatin accessibility and gene expression in progenitor cells, but did not survive correction for multiple comparisons for association with expression in neurons (Fig. 5c). Mediation analysis showed that this variant impacts *SLC26A7* gene expression through chromatin accessibility (Fig. 5d-e, posterior causal forward complete/causal forward partial: 0.028/0.966). Mutations in SLC26A7 protein were found in individuals with congenital hypothyroidism, though the role of this gene in brain structure is unclear [68–70]. Overall, here we proposed a cell-type-specific genetic causal path where regulation of *SLC26A7* gene impacts brain structure, possibly through thyroid metabolism, that can be experimentally validated in future studies.

**Figure 5.**
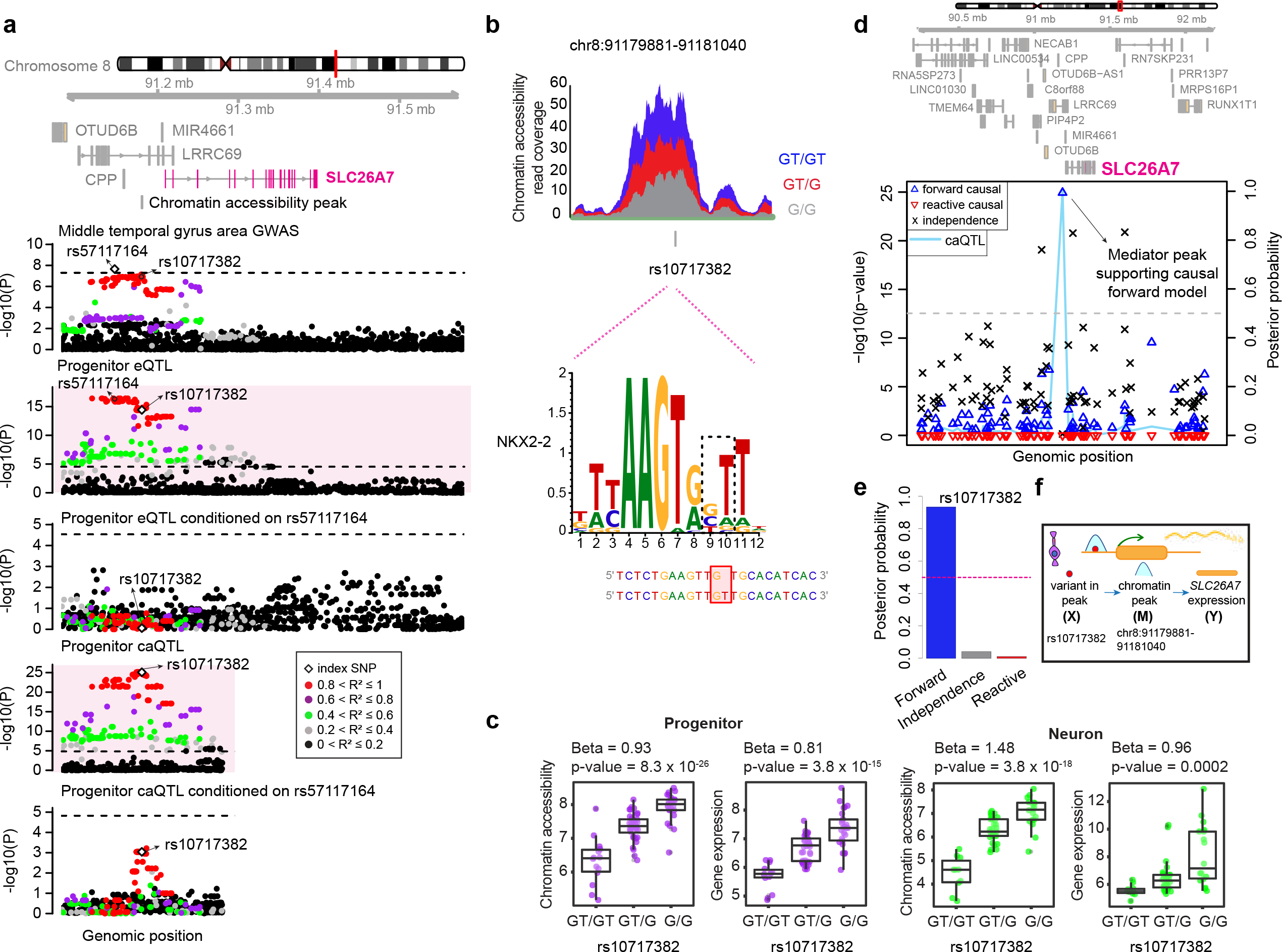
Colocalization of shared ca/eQTLs at *SLC26A7* locus with middle temporal gyrus area GWAS. **a)** Genomics tracks illustrating regional association of variants with *SLC26A7* gene expression, chromatin accessibility at its promoter, and middle temporal gyrus area GWAS. Colocalization of middle temporal gyrus area GWAS index SNP with shared ca/eQTL SNP was detected via conditional analysis. Data points were colored based on the pairwise LD r^2^ with the variant rs10717382 within the chromatin accessible region. **b)** Coverage plot illustrating ATAC-seq reads within the chromatin accessible region per genotype. Genomic position of the variant rs10717382 was shown along with the NKX2-2 TF motif disrupted by rs10717382. **c)** Phenotype (chromatin accessibility and *SLC26A7* expression) versus genotype boxplots per cell-type. **d)** Mediation scan plot overlaid with caQTL data for the causal forward model of epigenetic regulation of *SLC26A7* expression in progenitors indicated by lines in different colors with corresponding p-value on the left y-axis, and the posterior probability of each possible model showed with different shapes on the right y-axis. The location of the peak supporting causal forward model was highlighted. **e)** bmediatR posterior probability for causal forward, independence versus causal reactive models for the regulation of *SLC26A7* by chromatin accessibility. **f)** The cartoon illustrating causal forward model for the regulation of *SLC26A7* by chromatin accessibility.

We also utilized trans-eQTL mediation approaches to yield a deeper understanding of the mechanisms of how genes influence brain traits (Table S3). We identified a variant (rs10434445) that was significantly associated with *SRP72* expression via cis-regulation on chromosome 4 and was also a trans-eQTL for *TUBG2* on chromosome 17 in progenitors (Fig. 6a-b). These two genes were positively correlated (Fig. 6c), and mediation analysis showed that the genetic effect of the variant on *TUBG2* was mediated via *SRP72* gene expression (posterior probability = 0.813, Fig. 6d-e). Importantly, we previously found *TUBG2* as a TWAS significant gene for educational attainment (EA) from a GWAS in the same cell type [14] (Fig. 6f). *SRP72* encodes a signal recognition particle that is responsible for intracellular trafficking [71]. *TUBG2* was found to be highly expressed in the mouse brain and encoded protein functions in microtubule nucleation [72].

**Figure 6.**
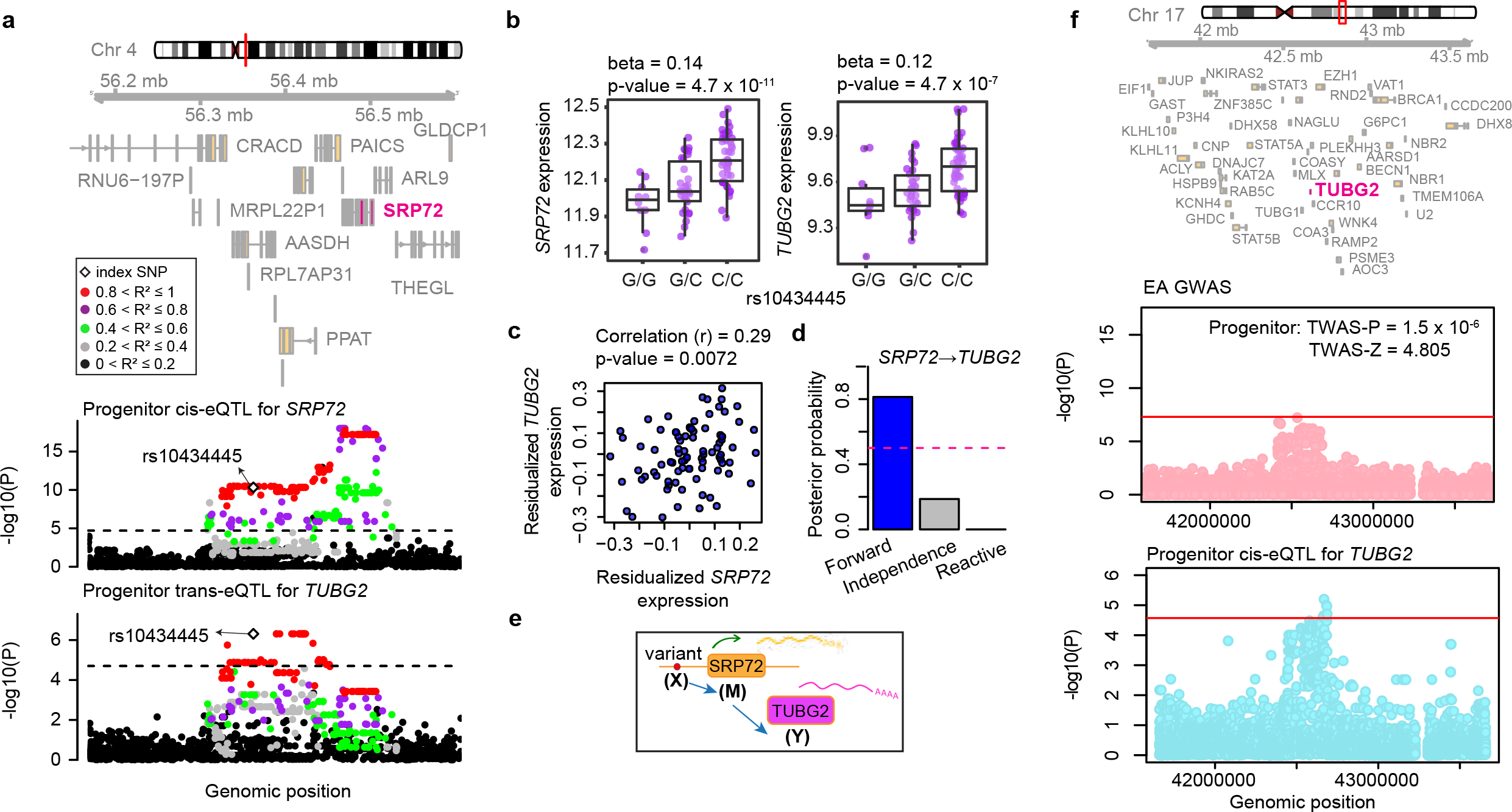
Trans-regulation *TUBG2* gene *via SRP72* gene. **a)** Genomics tracks illustrating association of variants cis-regulating *SRP72* upstream gene with the expression *SRP72*, and downstream gene *TUBG2*. Data points were colored based on the pairwise LD r^2^ with the rs10434445. **b)** Phenotype (*SRP72* and *TUBG2* expression) versus genotype (rs10434445) boxplots in progenitors. **c)** Correlation of *SRP72* and *TUBG2* expression. **d)** bmediatR posterior probability for causal forward, independence versus causal reactive models for the regulation of *TUBG2* by *SRP72*. **e)** The cartoon illustrating causal forward model for the regulation of *TUBG2* by *SRP72*. **f)** Regional association of variants with EA GWAS (upper, pink) and *TUBG2* gene (lower, light blue). EA TWAS p-value and z-score were shown for *TUBG2*.

## Discussion

Here, integrating multiple QTLs, we identified cell-type-specific causal gene regulatory mechanisms in a model system of the developing human brain. Our approach demonstrated several important features of gene regulation: (1) testing causality for shared QTLs between chromatin accessibility and gene expression, and between cis and trans-regulated genes provided insights into epigenomic and transcriptomic features of gene regulatory networks by linking variant to peak to gene and variant to gene to gene; (2) utilization of a cell-type-specific system allowed us to observe context-dependent causal networks that may have been masked by tissue heterogeneity in previous causal inference efforts on bulk tissue from the human brain; (3) applying a Bayesian strategy rather than traditional regression based mediation, we could evaluate models of genetic effects on gene expression via chromatin accessibility in terms of posterior probability of fully, partially, or independent mediation.

Different molecular assays require unique experimental procedures resulting in different levels of technical noise across molecular phenotypes, which may bias estimates of mediated effect [23, 73]. Importantly, based on simulations, we demonstrated that reliable measurements are required to make accurate conclusions about causal mechanisms after data integration. We attempted to eliminate false positive causal relationships by implementing an algorithm that detects a threshold ICC value. We observed that forward models can be identified as reactive at the low ICC values; whereas reactive to forward model flipping is very unlikely to occur. Several adjustments including larger sample sizes, more donors with replicates, and a higher read coverage for sequencing data might reduce the measurement error. Moreover, despite our attempt to avoid potential false positive reactive models via this strategy, we were not able to identify any biologically meaningful examples of reactive candidates. This observation may be interpreted in light of the fact that (1) observation of a reactive model, chromatin accessibility mediated by gene expression, is an unlikely biological mechanism in this dataset, and (2) small sample size and measurement error may obscure distinguishing partial forward and partial reactive, as observed by the authors of the *bmediatR* method in simulation analyses [22]. We expect that increasing the statistical power of QTLs using a larger number of samples will be helpful to determine how often, if at all, reactive regulatory models exist.

We explored a variant co-localized with an index SNP for the middle temporal gyrus area upstream of *SLC26A7* gene, which alters chromatin accessibility at the region leading to changes in gene expression. This observation directed us to propose a comprehensive mechanism to interpret how a GWAS locus alters complex brain structure. Despite the low number of QTLs that colocalized with any brain-relevant GWAS loci, we detected cis-regulatory elements mediating the expression of genes whose haploinsufficiency is associated with rare neurodevelopmental conditions. For instance, we found that a genetically altered chromatin accessible region upstream of the *CHL1* gene mediated its expression. Heterozygous loss of the *CHL1* gene has been observed in individuals with cognitive delays [54, 55], though a definitive statistical association has not yet been conducted. Upon validation by cellular assays, this regulatory region could be an appealing candidate upstream enhancer to modulate endogenous gene expression for CRISPR-based targeted therapeutic approaches [74–77]. In addition to epigenetic regulations, we identified an upstream cellular target *SRP72* gene regulated by cis-regulatory elements for EA TWAS gene *TUBG2*. Using the genetic variation within this region as an instrumental variable serving as a natural multifactorial perturbation [78], we could build a causal relationship between two genes that would not be possible to infer by merely considering their co-expression. Exploring the other genes regulating the disease-associated gene can help to gain a comprehensive understanding of the underlying mechanisms of human intelligence and to find novel disease-relevant intracellular pathways.

Although our sample size was comparable with other cell-based QTL studies [79–82], increasing the number of donors will likely lead to a higher number of shared QTLs to be tested in mediation analysis. Also, since the detection of some QTLs is dependent on the presence of stimuli, this may prevent simultaneous observation of changes in chromatin accessibility and gene expression [83, 84]. To this end, further studies using more context-dependent conditions can be complementary to our findings with an objective to examine causal networks that connect risk variants associated with brain-relevant traits to cellular function.

## Conclusion

In this study, we identified epigenetic and trans-regulated causal pathways for the underlying mechanism of gene expression, employing a cell-type specific system representing a critical period of human cortical development. Leveraging these causal networks with brain-relevant GWASs, we proposed potential molecular functions for trait associated variants, which represent novel candidates for mechanistic studies aiming to understand inter-individual differences in neurodevelopmental traits.

## Methods

### Establishment of primary human neural progenitor cells (phNPCs)

We generated phNPCs and differentiated them into neurons following the same cell culture procedure as described in our previous work [13,14,85].

### Generation of cell-type-specific ATAC-seq and RNA-seq data

We prepared ATAC-seq libraries, sequenced them using Illumina HiSeq2500 or MiSeq platforms with 50 bp paired-end sequencing and aligned them to the human genome (GRCh38/hg38) by reducing mapping bias via WASP method after quality control as described previously [13]. We generated RNA-seq libraries, performed sequencing with NovaSeq S2 flow cell using 150 bp paired-end sequencing, and also mapped them to the human genome (GRCh38/hg38) after quality control as previously described [14].

### Genotype processing and imputation

Genotyping was performed using Illumina HumanOmni2.5 or HumanOmni2.5Exome platforms. If variants showed variant missing genotype rate > 5% (--geno 0.05), deviations from Hardy-Weinberg equilibrium at p <1×10^−6^ (--hwe 10^−6^), and minor allele frequency < 1% (--maf 0.01), we filtered them out. We also excluded samples if they had missing genotype rate > 10% (--mind 0.10) as described previously [13, 14]. For imputation, we used 1000 Genomes Project Phase 3 reference panel for multiple ancestries [86] using Minimac4 software [87] by retaining variants with missing genotype rate lower than 0.05, Hardy-Weinberg equilibrium p-value greater than 1 × 10^−6^, minor allele frequency (MAF) bigger than 1% and imputation R^2^ greater than 0.3 as described previously [13, 14].

### Intraclass correlation within chromatin accessibility and gene expression data

To quantify cell culture-induced noise, we cultured 11 and 5 donors in progenitors and neurons to prepare ATAC-seq libraries and 11 and 9 donors in progenitors and neurons to prepare RNA-seq libraries multiple times during the course of the experiment. We calculated the intraclass correlation coefficient (ICC) of gene expression and chromatin accessibility between libraries from the same donors. For neurons, we used gene expression values after batch correction with the limma R package for the sorter type, as described above. We performed an unpaired two-sided t-test for statistical assessment of the mean difference between these two categories (Fig. S4a).

### Cell-type-specific eQTL and caQTL mapping to test causal forward model

To find the candidate SNP-chromatin accessibility-gene candidates to be tested for causal forward model against independence, we tested association of variants within chromatin accessible region with the chromatin accessibility in these regions, and with the genes if the variants were within +/-1MB gene TSS window. Using the same models as previously established for caQTLs and eQTLs for each cell-type, we re-performed QTL analyses including the donors that have both ATAC-seq and RNA-seq phenotypes (N_donor_ = 75 in progenitors and N_donor_ = 57 in neurons) [13, 14] as follows:

Progenitor caQTL: chromatin accessibility ∼ SNP + 10 MDS of global genotype + kinship matrix + 8 PCs of global gene expression

Neuron caQTL: chromatin accessibility ∼ SNP + 10 MDS of global genotype + kinship matrix + FACS sorter + 7 PCs of global gene expression

Progenitor eQTL: expression ∼ SNP + 10 MDS of global genotype + kinship matrix + 10 PCs of global gene expression

Neuron eQTL: expression ∼ SNP + 10 MDS of global genotype + kinship matrix + FACS sorter + 12 PCs of global gene expression

We applied the same hierarchical multiple testing correction strategy as previously by computing a global eigenMT-FDR p-value after local adjustment per eGene, and defined significance eQTLs at 5% eigenMT-FDR [13, 14]. For caQTLs, we adjusted association p-values via only the FDR method given that there were a few chromatin accessible regions with multiple variants, and retained the associations with lower than 5% FDR.

### Cell-type specific eQTL and caQTL mapping to test causal reactive model

To find the candidate SNP-chromatin accessibility-gene triplets to be tested for causal reactive model against independence, we applied the following strategy: (1) we detected (i) eGenes (53 eGenes in progenitors and 12 eGenes in neurons) that encode transcription factors with known motifs [88] by calculating eigenMT-FDR threshold p-value by using only eGenes encoding TFs, and (ii) variants cis-regulating these eGenes at this significant threshold; (2) we searched chromatin accessible regions throughout the genome that harbor 80% matching sequence of the binding motifs [88] of these TF eGenes via TFBSTools [89]; and (3) we performed trans-caQTL analysis by using variants from step 1 and chromatin accessibility at the regions from step 2 with the same caQTL models described for the causal forward model. We defined significant trans-caQTLs at 5% FDR.

### Cell-type specific trans eQTL mapping

We tested associations of variants that were cis-regulating at least one gene in our previous analysis (N_donor_ = 85 in progenitors and N_donor_ = 74 in neurons) [14] with all the genes in the genome expressed in our dataset. We included variant-gene pairs, if the distance between variant and TSS of the gene was larger than 1MB or they were on different chromosomes. We used the following models:

Progenitor eQTL: expression ∼ distal SNP + 10 MDS of global genotype + kinship matrix + 10 PCs of global gene expression

Neuron eQTL: expression ∼ distal SNP + 10 MDS of global genotype + kinship matrix + FACS sorter + 12 PCs of global gene expression

We retained the associations with a nominal p-value lower than 1 × 10^−6^ following the previous trans-eQTL strategies [46, 47], and defined X (single variant)-M (cis-regulated or upstream gene expression)-Y (trans-regulated or downstream gene expression) triplets. We found that the linear mixed-effects model used to identify eQTLs can be susceptible to instability during fitting with large numbers of covariates. We removed 5 and 1 cis-eGenes in progenitors and neurons that had large differences in beta values prior to and after the inclusion of covariates.

### Bayesian approach for mediation analysis

For mediation of gene expression through chromatin accessibility: for each X-M-Y triplets where X is a genetic variant (encoded as -1,0,1) within a chromatin accessibility region that is significantly associated with both chromatin accessibility and gene expression; M is the chromatin accessibility and Y is the gene expression, we ran mediation analysis applying bmediatR [22] using the covariates for caQTL and eQTL data corresponding to population structure and technical factors. We applied the default setting for hyperparameters representing the effect sizes to φ^2^ = (1, 1, 1) relationships between X and M, M and Y, and X and Y, respectively, assuming them to be equal *a priori* for each, since we observed that φ^2^ = (1, 1, 1) as one of the values maximizing the sum of marginal log likelihoods (Fig. S1c). We used the non-informative default priors for the scaling parameters (κ, λ) = (0.001, 0.001) and fixed effect coefficients, both the intercept and covariates τ = (1000, 1000). This analysis calculated the posterior distribution of θ, which denotes the edges of the DAG relating X-MY-by multiplying a joint likelihood for Y and M with a prior distribution for θ as p(θ|y,m) ∝ p(y,m|θ)p(θ). We defined the relationship as causal forward if the sum of the posterior probabilities of complete and partial mediation was higher than 0.5, and as reactive if the sum of the posterior probabilities of complete reactive and partial mediation models was higher than 0.5.

To limit potential false positive (FP) causal relationships that may result from imbalanced measurement error across M and Y, we simulated normally distributed random variables introducing variable error added to M (M’) and Y (Y’). We then detected false positive causal reactive and forward mediation, respectively, using percent of variance of M explained by X (PVE_A) and the percent of variance of Y explained by M on the odds scale (PVE_B) per X-M-Y triplet from the real data. We performed analysis separately for each scenario of model flipping: (1) from forward to reactive to detect FP reactives, and (2) from reactive to forward to detect FP forwards. After running mediation analysis, we fitted the posterior probabilities supporting forward and reactive models against ICC values changed upon application of the error term via local polynomial regression. Following the fitting, we attempted to define an ICC threshold at which the model flips from forward to reactive or from reactive to forward. To this end, we calculated the distance between the two curves corresponding to two models and defined the ICC value that minimized the distance if the slope of the wrong model is positive and the slope of the correct model is negative relative to ICC values bigger than this threshold (Fig. S4c). We performed this simulation 10 times and averaged ICC threshold values per X-M-Y triplet whose corresponding PVE_A and PVE_B values were used during the simulation. We filtered out X-M-Y triplets with ICC for chromatin accessibility was lower than the threshold ICC value for the results supporting the reactive model, respectively. Additionally, we retained only X-M-Y triplets with positive ICC values for chromatin accessibility and gene expression or both models. We detected that model flipping from reactive to forward was only possible when both PVE_A and PVE_B values were bigger than 0.9 (data not shown). Since we did not have such large PVE values for X-M-Y triplets tested in our study, we did not need to apply this algorithm to limit potential false-positive forwards.

For mediation of gene expression through the expression of a distal gene, each X-M-Y triplet where X is a genetic variant that is significantly associated with the expression of both genes; M is the gene cis-regulated by X and Y is the gene trans-regulated by X, similarly, we applied bmediatR [22]. We used the covariates for eQTL data corresponding to population structure and technical factors with the same hyperparameters described for mediation through caQTL above. Given that the reactive model was not biologically interpretable for trans-regulation, we only considered the forward direction for causality, and we considered the relationship as causal forward if the sum of the posterior probabilities of complete and partial mediation were higher than 0.5.

### Regression-based approach for mediation analysis

We additionally applied a traditional mediation approach with regression analysis to detect the mediation of genetic effects on gene expression via chromatin accessibility. For each cell type, we established the linear models where ZY and ZM are the covariate matrices used for eQTL and caQTL analyses, Mresidualized is chromatin accessibility residualized by ZM and Yresidualized is gene expression residualized by ZY and tested alternative hypothesis H1 below:

For causal forward model:

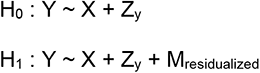

For causal reactive model:

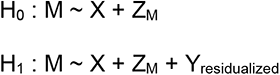

We performed multiple testing applying a mediation scan strategy as previously described [44]. For a causal reactive/forward model, per each peak–SNP pairs/gene–SNP pairs, we randomly selected 50 genes/chromatin accessibility regions from another chromosome to condition on. Then after permuting the mediator 1000 times for the sample index, we selected the maximum conditional p-value (the least significant) from each permutation and fit a generalized extreme value distribution (GEV). Finally, we calculated the FWER-controlled mediation p-values using the cumulative density function of the GEV and set a significance threshold of 0.05.

To evaluate a forward model with a high partial mediation probability for *CHL1* gene regulation (Fig. S2c), we applied a similar strategy. We tested if there was any significant association between residualized *CHL1* expression by both technical covariates and center peak chromatin accessibility (that was also residualized by technical covariates), and peak 1 and 2 chromatin accessibilities, separately (that were also residualized by technical covariates).

### Regression-based mediation analysis with MOSTWAS

We performed MOSTWAS analysis as a regression-based mediation approach to detect trans-eQTLs mediated by cis-eQTLs by following the procedure described previously [47]. In brief, for each SNP-cis gene (upstream)-trans gene (downstream) triplets, we estimated total indirect mediation effect of the distal SNPs (cis SNPs for the upstream gene) on the downstream gene of interest using a permutation approach (nperm = 1000), and the resulting permutation p-values were adjusted to control the FDR. We considered the downstream genes with permutation p-values lower than 10% FDR and with heritability p-value lower than 0.05 as mediated by upstream genes.

### GWAS co-localization analysis

To find eQTLs colocalized with index GWAS loci, we performed LD-thresholded colocalization analysis for each cell type separately [90]. We used summary statistics from GWAS for schizophrenia (SCZ) [2], major depression disorder (MDD) [91], educational attainment (EA) [6] and cortical thickness and surface area from ENIGMA project [5], UKBB [4], bipolar disorder (BP)[92], neuroticism[93], IQ[94], cognitive performance (CP) [6], attention-deficit/hyperactivity disorder (ADHD) [95], Alzheimer’s disease (AD)[96], Parkinson’s disease (PD) [97]. We defined index GWAS SNPs where two LD-independent GWAS signals so as to have pairwise LD r^2^ < 0.2 based on LD matrix computed with European population of 1000 Genomes (1000G European phase 3) at genome-wide significant threshold p-value (5×10^−8^). Then, we found (1) two highly correlated variants (pairwise r^2^ between GWAS and QTL index variant was higher than 0.8 based on either our study or European population), and (2) if the gene expression/chromatin accessibility was no longer significantly associated with QTL index variant conditioning on GWAS index variant, these two loci were co-localized.

### Cross-mappability of genes

We used a pre-computed multi-mapping score from a previous study [59]. For each upstream-downstream gene pair, we used symmetric cross-mappability between upstream gene (gene A) and downstream gene (gene B) that was calculated as (crossmap(A,B) + crossmap(B,A))/2 [59]. We discarded upstream-downstream gene pairs if the cross-mappability scores between them were higher than 5 at the log2 scale.

### Transcription factor motif analysis

We used motif breaker R to detect the disruption of the transcription motif binding site where there was a variant within a chromatin accessibility peak [57]. To detect transcription motifs within gene promoters, we used TFBStools [89] with 80% minimum matching score by searching a target sequence within +/-500 bp window from gene TSS.

## Declarations

### Ethics approval and consent to participate

This study was reviewed by the UNC Office of Human Research Ethics (16-0054) and was determined to be not human subjects research so does not require IRB approval.

### Consent for publication

Not applicable.

### Availability of data and materials

Codes are available here https://bitbucket.org/steinlabunc/pathqtl/src/master/

### Authors’ contributions

JLS and MIL conceived and supervised the study. MIL and JLS provided funding. NA managed the integration of the datasets, interpreted results and performed caQTL, cis and trans eQTL, mediation, co-localization, enrichment, transcription factor binding motif analyses, and conducted simulations. DL performed pre-processing of genotyping data and ATAC-seq data. WLC and GRK aided in *bmediatR* methodology. JLS, MIL and NA wrote the manuscript. All authors commented on and approved the final version of the manuscript.

## Acknowledgments

The following core facilities were utilized for this project: UNC Neuroscience Center Microscopy Core (P30NS045892), UNC Mammalian Genotyping Core, CGIBD Advanced Analytics Core (NIH grant P30 DK034987), UNC Flow Cytometry Core Facility, UNC Vector Core, UNC Research Computing. Additional core facilities utilized for this project were: UCLA CFAR (5P30 AI028697), and the UCLA Neuroscience Genomics Core. We thank Dr. William Valdar for helpful comments regarding Bayesian mediation analysis.

## Supplemental figure legends

**Figure S1.**
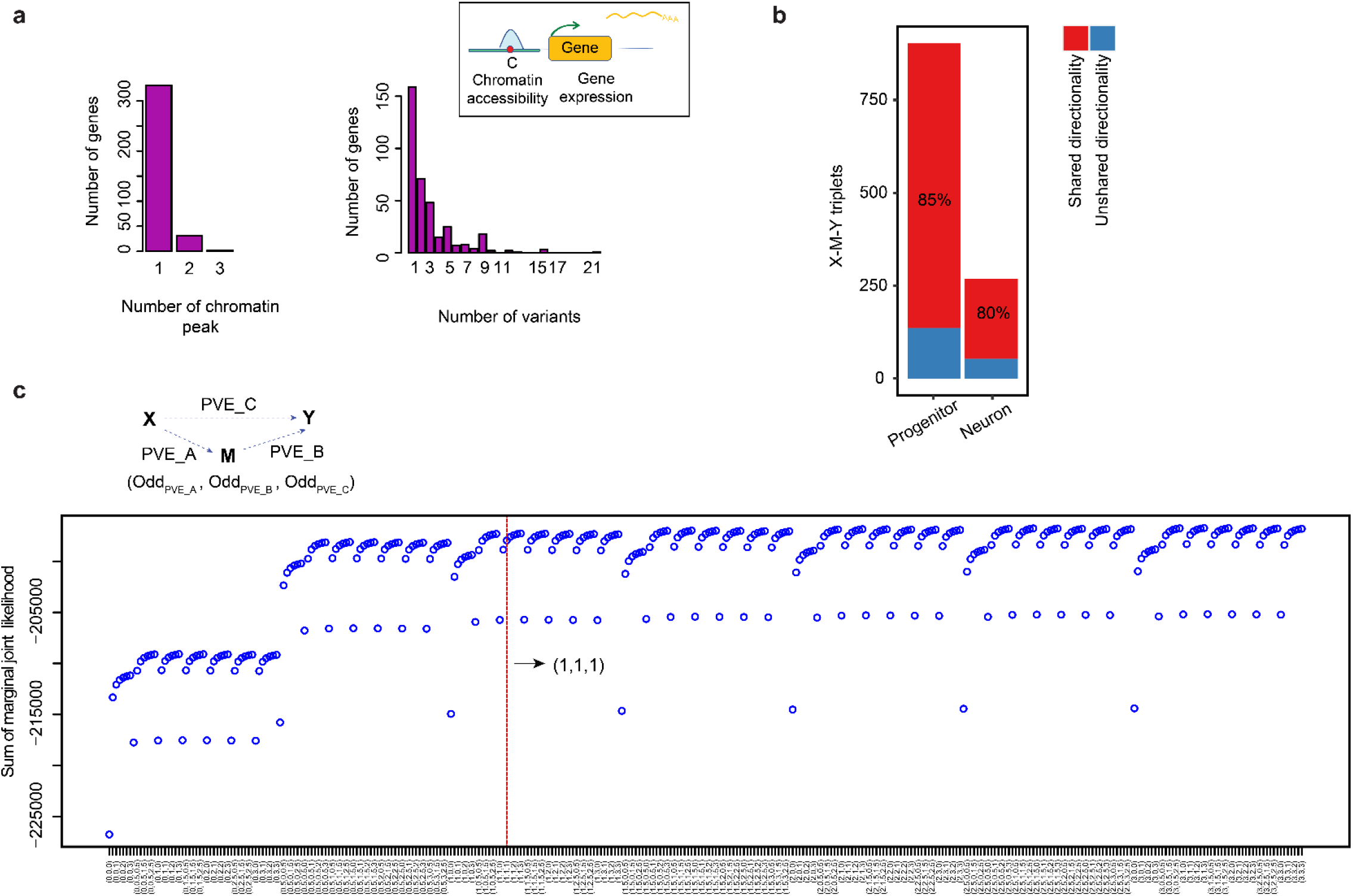
Colocalized ca/eQTLs and hyperparameter selection for bmediatR. **a)** The number of variants, chromatin accessible regions and gene expression tested as candidate X-M-Y triplets. **b)** Proportion of shared and unshared candidate X (genetic variant)-M (chromatin accessibility)-Y (gene expression) triplets based on directionality of the genetic effect for each cell-type. **c)** Sum of marginal joint likelihood at different PVE_A, PVE_B and PVE_C hyperparameters at odds scale.

**Figure S2.**
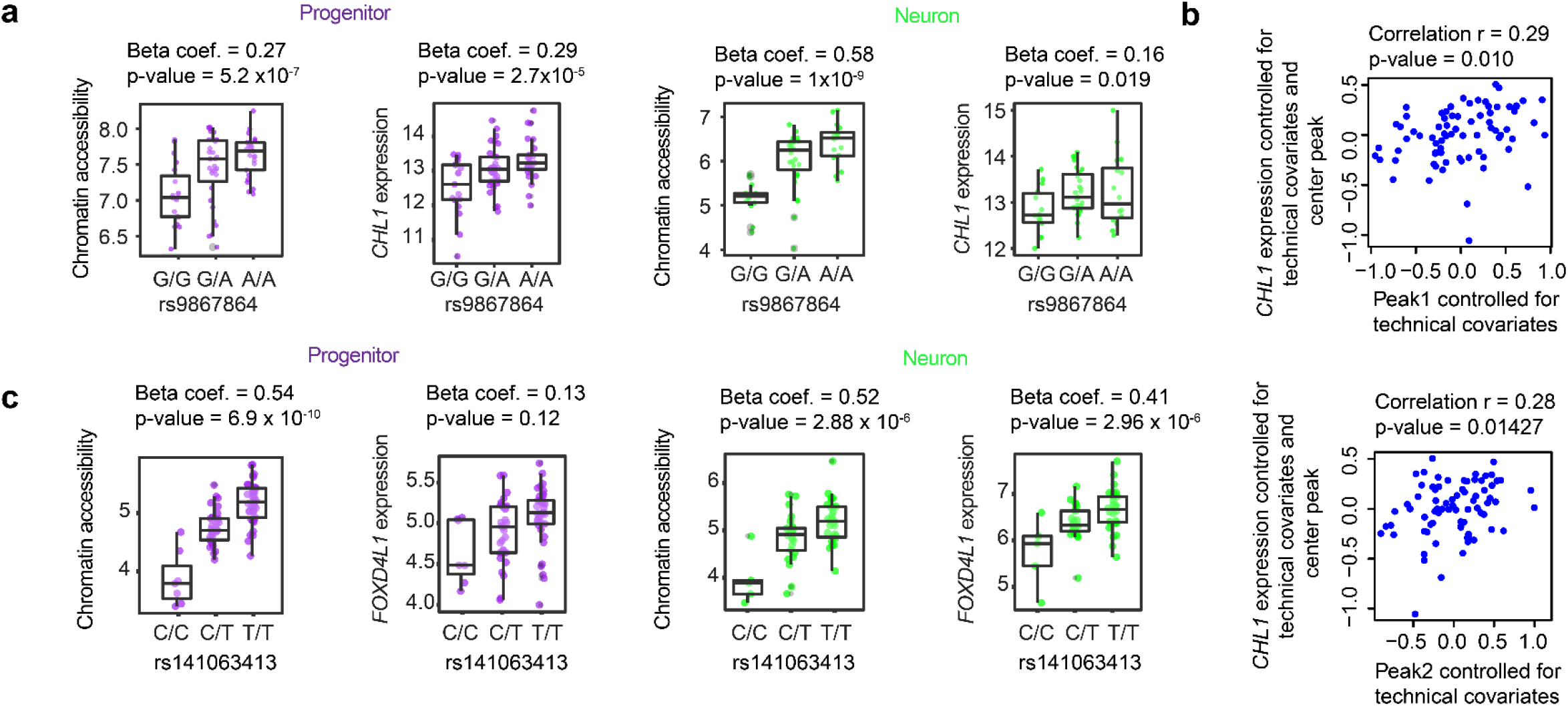
ca/eQTLs at *CHL1* and *FOXD4L1* loci for each cell type. **a)** Phenotype (chromatin accessibility and *CHL1* expression) versus genotype boxplots per cell-type. **b)** Association of the residualized *CHL1* expression by technical covariates and chromatin accessibility at center peak residualized by technical covariates with the chromatin accessibility at peak1 (upper plot) or peak2 (lower plot) residualized by technical covariates **c)** Phenotype (chromatin accessibility and *FOXD4L1* expression) versus genotype boxplots per cell-type.

**Figure S3.**
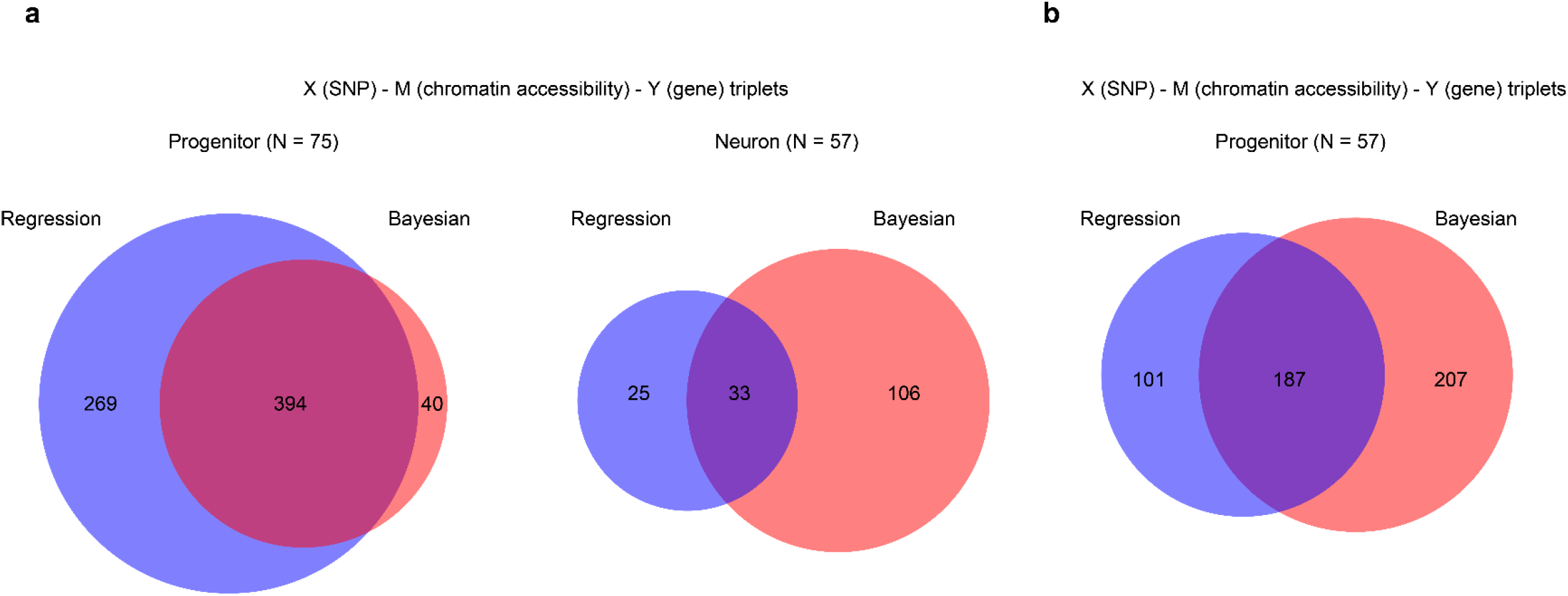
Comparison of bayesian and regression based mediation analyses. **a)** Comparison of X-M-Y triplets supporting causal forward model detected by regression based versus bmediatR method in progenitors (N_donor_ = 75) and neurons (N_donor_ = 57). **b)** Comparison of X-M-Y triplets supporting causal forward model detected by regression based versus bmediatR method in progenitors after they were downsampled (N_donor_ = 57).

**Figure S4.**
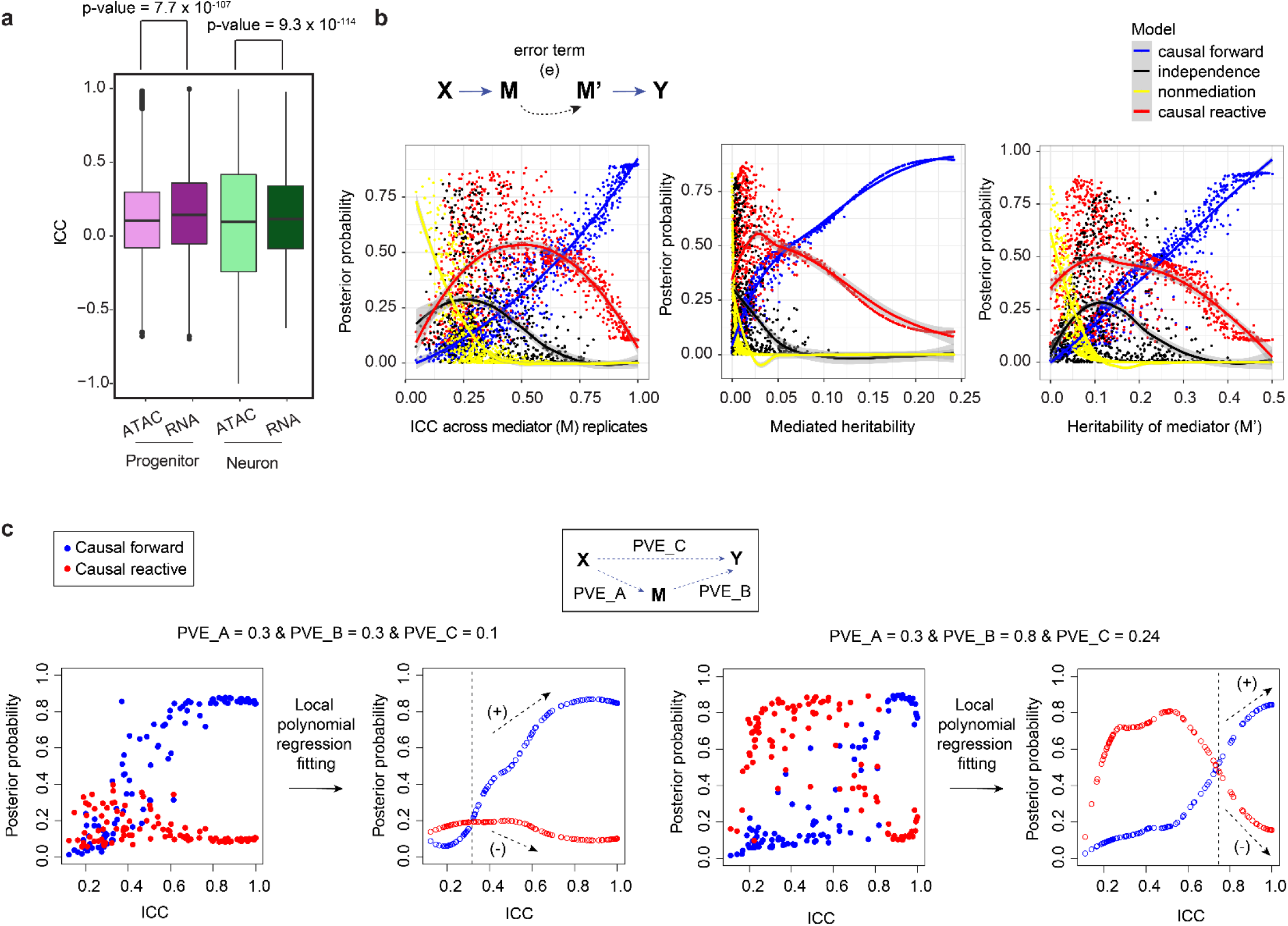
Measurement error differences between ATAC-seq and RNA-seq and detection of false positive reactive models. **a)** Intraclass correlation coefficient (ICC) for ATAC-seq measured peaks and RNA-seq measured genes in progenitors and in neurons. Unpaired t-test p-values were shown. **b)** Simulation analysis for model flipping from causal forward to causal reactive given the error term on mediator (M). The impact of ICC, mediated heritability and heritability of mediator values on model flipping. Posterior probability of each model was indicated by different colored lines. **c)** Depiction of the algorithm used to eliminate false positive reactive results at a low threshold ICC value.

**Figure S5.**
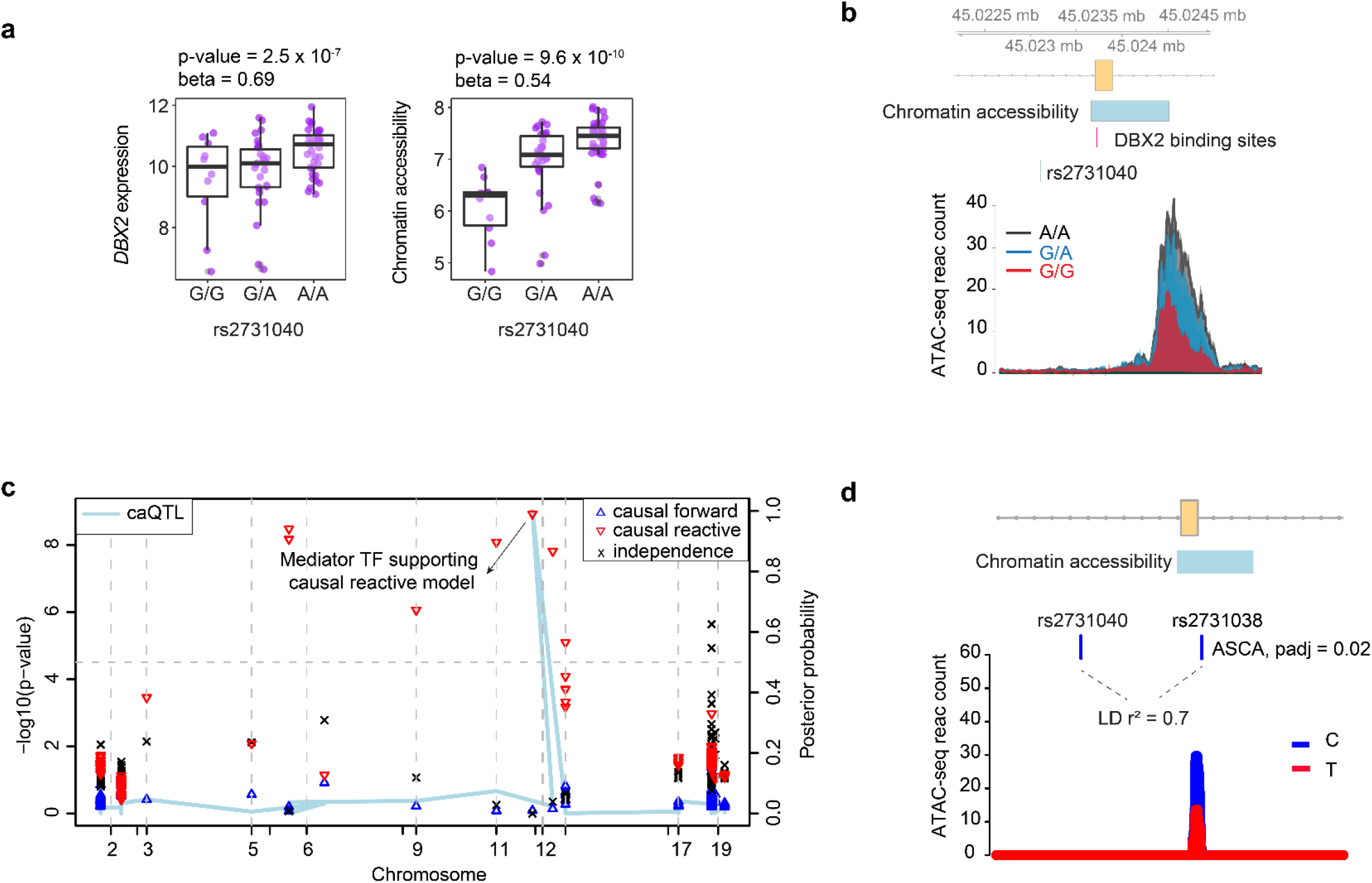
Evaluation of causal reactive model at *DBX2* locus. **a)** Genotype versus phenotype boxplots for *DBX2* expression and chromatin accessibility in progenitors. **b)** Location of chromatin accessibility within *DBX2* gene body, and coverage plot for chromatin accessibility across genotypes. **c)** Mediation scan plot illustrating causal reactive model whereby only *DBX2* gene expression leads to chromatin accessibility, but not any other genes encoding TFs with matching motifs within chromatin accessible region. **d)** Another variant, rs2731038, within the chromatin accessible region that was in LD with rs2731040 (r² = 0.7), showed allele-specific-chromatin accessibility (ASCA). Padj: Adjusted p-value after ASCA analysis.

**Figure S6.**
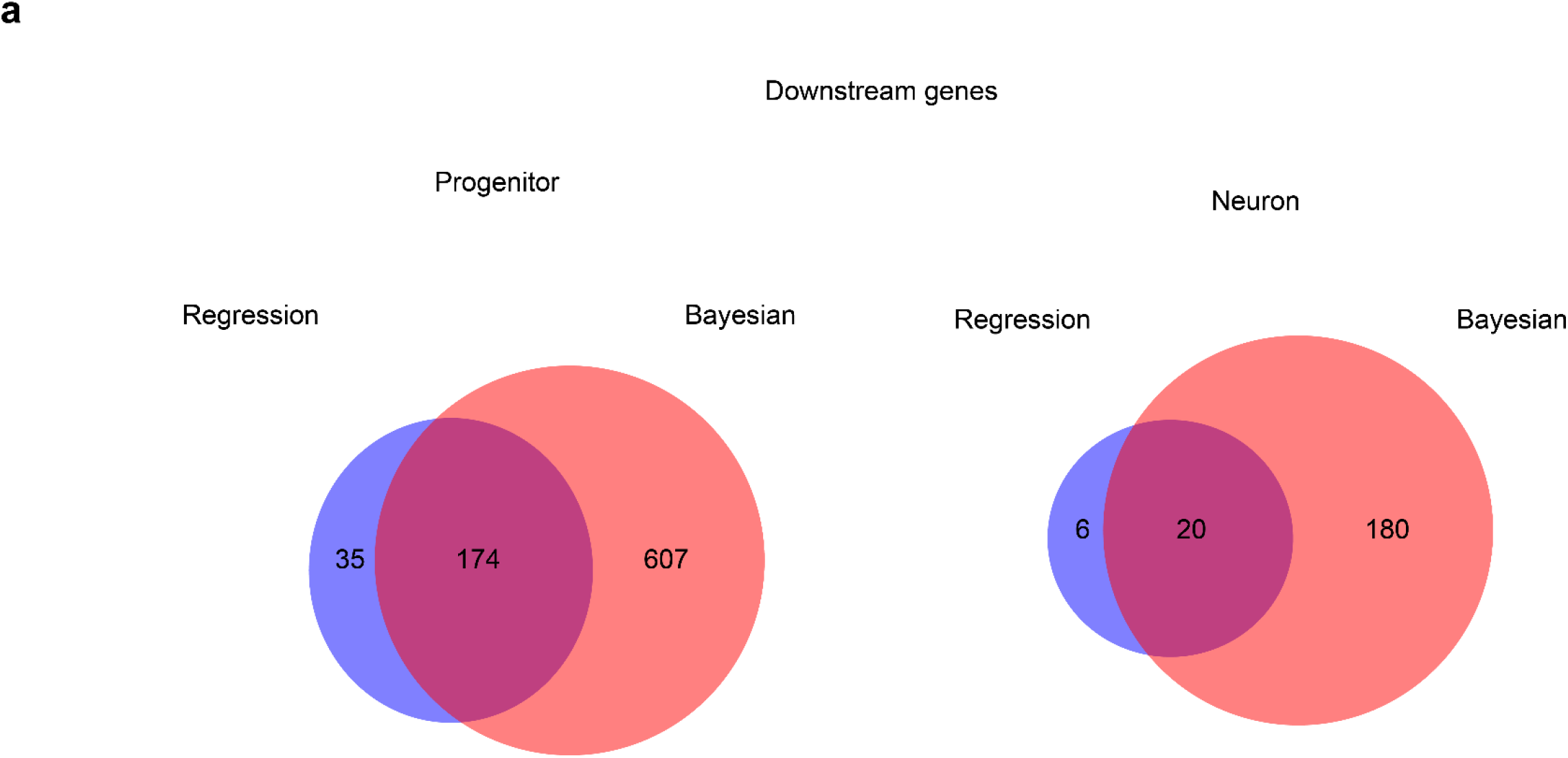
Comparison of bayesian and regression based mediation analyses. **a)** Comparison of X-M-Y triplets supporting causal forward model detected by regression based versus bmediatR method for gene mediated by trans-regulation.

## Supplementary tables

**Table S1:**Causal forward and reactive mediation results for ca/eQTL colocalizations.

Sheet 1: Causal forward: snp: variant tested; gene.beta: beta estimate of the eQTL; gene.pval: p-value for eQTL; gene: ensemblID for the gene of interest; peak: chromatin accessibility peak defined as chromosome, peak start and peak end peak.beta: beta estimate of the caQTL; peak.pval: p-value for caQTL; chr: chromosome where the variant is located; icc_gene: ICC value for gene; icc_peak: ICC_value for chromatin accessibility; complete.med: posterior probability of complete mediation in forward direction; partial.med: posterior probability of partial mediation in forward direction; co.local: posterior probability of independence; complete.med.reac: posterior probability of complete mediation in reactive direction; partial.med.reac: posterior probability of partial mediation in reactive direction; symbol: gene symbol; A1: effect allele; cell_type: cell-type used for analysis.

Sheet 2: Causal reactive: snp: variant tested; gene.beta: beta estimate of the eQTL; gene.pval: p-value for eQTL; gene: ensemblID for the gene of interest; peak: chromatin accessibility peak defined as chromosome, peak start and peak end peak.beta: beta estimate of the caQTL; peak.pval: p-value for caQTL; chr_snp: chromosome where the variant is located; chr_peak: chromosome where the peak is located; icc_gene: ICC value for gene; icc_peak: ICC_value for chromatin accessibility; complete.med: posterior probability of complete mediation in forward direction; partial.med: posterior probability of partial mediation in forward direction; co.local: posterior probability of independence; complete.med.reac: posterior probability of complete mediation in reactive direction; partial.med.reac: posterior probability of partial mediation in reactive direction; symbol: gene symbol; A1: effect allele; cell_type: cell-type used for analysis; icc_thres: ICC threshold value flipping the model.

**Table S2:**Causal forward and reactive mediation results for cis/trans eQTL colocalizations.

snp: variant tested; cis_beta: beta estimate of the cis-eQTL; cis_pvalue: p-value for cis-eQTL; cis_gene: ensemblID for the upstream gene of interest; trans_beta: beta estimate of the trans-eQTL; trans_pval: p-value for trans-eQTL; cis_chr: chromosome where the variant and upstream gene are located; trans_chr: chromosome where the downstream gene is located; complete.med: posterior probability of complete mediation in forward direction; partial.med: posterior probability of partial mediation in forward direction; co.local: posterior probability of independence; complete.med.reac: posterior probability of complete mediation in reactive direction; partial.med.reac: posterior probability of partial mediation in reactive direction; symbol: gene symbol; A1: effect allele; cell_type: cell-type used for analysis.

**Table S3:** Colocalization of ca/e/transSNPs with GWAS and TWAS genes

Sheet 1: Cell-type-specific ca/eQTL colocalizations with GWAS: snp: variant tested; gene.beta: beta estimate of the eQTL; gene.pval: p-value for eQTL; gene: ensemblID for the gene of interest; peak: chromatin accessibility peak defined as chromosome, peak start and peak end peak.beta: beta estimate of the caQTL; peak.pval: p-value for caQTL; chr: chromosome where the variant is located; icc_gene: ICC value for gene; icc_peak: ICC_value for chromatin accessibility; complete.med: posterior probability of complete mediation in forward direction; partial.med: posterior probability of partial mediation in forward direction; co.local: posterior probability of independence; complete.med.reac: posterior probability of complete mediation in reactive direction; partial.med.reac: posterior probability of partial mediation in reactive direction; symbol: gene symbol; condBeta: conditional beta estimate; condP: conditional p-value; coloc.type: if caQTL or eQTL was colocalized; gwas_SNP: GWAS index SNP; A1: effect allele; trait: GWAS trait; cell_type: cell-type used for analysis.

Sheet 2: Cell-type-specific cis/trans eQTL colocalizations with GWAS: snp: variant tested; cis_beta: beta estimate of the cis-eQTL; cis_pvalue: nominal p-value for cis-eQTL; cis_gene: ensemblID for the upstream gene of interest; trans_beta: beta estimate of the trans-eQTL; trans_pval: nominal p-value for trans-eQTL; cis_chr: chromosome where the variant and upstream gene are located; trans_chr: chromosome where the downstream gene is located; complete.med: posterior probability of complete mediation in forward direction; partial.med: posterior probability of partial mediation in forward direction; co.local: posterior probability of independence; complete.med.reac: posterior probability of complete mediation in reactive direction; partial.med.reac: posterior probability of partial mediation in reactive direction; condBeta: conditional beta estimate; condP: conditional p-value; gwas_SNP: GWAS index SNP; A1: effect allele; trait: GWAS trait (with IDP number for UKBB brain related traits); cell_type: cell-type used for analysis. Sheet 3: Cell-type-specific TWAS genes (upstream or downstream from trans-eQTL mediation analysis). Output from FUSION [98] and our previous study [14]: ID: the gene ensemblID or intron id; CHR: the chromosome number; HSQ: heritability; BEST.GWAS.ID: GWAS SNP in the locus with the most significant association; BEST.GWAS.Z: z-score of the best GWAS SNP; EQTL.ID: the best eQTL in the locus; EQTL.R2: the cross-validation R^2^ of the best eQTL in the locus; EQTL.Z: the z-score of the best eQTL in the locus; EQTL.GWAS.Z: the GWAS Z-score for this eQTL; NSNP is the number of SNPs in the locus; NWGT is the number of snps with non-zero weights; MODEL is the best performing model; MODELCV.R2 is the the cross-validation R^2^ of the best performing model; MODELCV.PV: the p-value from the cross-validation of the best performing model; TWAS.Z is the TWAS z-score; TWAS.P: TWAS p-value; trait is the GWAS trait; pop: the population used to estimate LD; joint_independent is the status if a gene/intron jointly independent; trait is the GWAS; gene_type: if the gene is upstream or downstream gene; cell_type: cell type for observation.

